# The TEAIV model: Extending the standard TEIV model to account for viral budding ramp up

**DOI:** 10.1101/2025.11.07.687137

**Authors:** S. Gholami, C.S. Korosec, Q. Su, M.I. Betti, J.M. Conway, I.R. Moyles, J. Watmough, J.M. Heffernan

**Affiliations:** Modelling Infection and Immunity Lab, Mathematics and Statistics, York University, Toronto, ON, Canada; Centre for Disease Modelling, Mathematics and Statistics, York University, Toronto, ON, Canada; Department of Mathematics and Computer Science, Mount Allison University, Sackville, NB, Canada; Department of Mathematics, Pennsylvania State University, University Park, Pennsylvania, United States of America; Department of Mathematics and Statistics, University of New Brunswick, 3 Bailey Dr, Fredericton, NB, Canada

## Abstract

Understanding how viruses replicate and spread within a host is fundamental to predicting disease progression, timing, and dosage of effective therapeutic and prophylactic interventions. A target-cell limited approach is often used to model within-host viral kinetics to characterize disease infection dynamics. The standard target-cell limited model, the TEIV model, has been instrumental for understanding SARS-CoV-2 within-host viral kinetics, however, its core assumptions of instantaneous viral budding and exponentially distributed cell lifetimes oversimplify fundamental biological processes, potentially limiting predictive power. In this work, we consider a novel model approach, the TEAIV model: an extension to the TEIV model that considers an infected partially-productive cell state that ramps up through *n* stages to a fully productive state. We fit the TEAIV model to various SARS-CoV-2 viral load datasets, separated by hospitalized (severe infection) and non-hospitalized (mild infection) individual data, consider multiple different ramping functions, and compare to standard TEIV model fits. We further compare TEAIV and TEIV model fits with and without infected cell death incorporated in the eclipse (E) and partially infected (A) compartments and investigate best-fit frameworks up to 10 A states. We find that linear budding ramp-up dynamics minimizes the BIC (with ΔBIC *>* 2) across all model formulations when the total number of eclipse and budding compartments exceeds three. Furthermore, we find that the inclusion or exclusion of cell death applied to eclipse or ramping compartments does not substantially affect this result. Finally, we analytically consider the most general TEAIV model and discuss future modelling considerations. Our results demonstrate that accounting for non-instantaneous viral budding provides a better fit to SARS-CoV-2 viral load data and establishes a foundation for more mechanistically informed within-host models.

## 1 Introduction

Mechanistic mathematical models of viral dynamics are essential in understanding the complex interplay between virus replication and host immune responses [1–8]. Mechanistic models provide quantitative insights into viral kinetics [9, 10], guide understanding of prophylactic [11–17] and anti-viral therapeutic in-host interactions [18– 21], bridge observations from within-host to population-scale transmission dynamics [22], and provide an intuitive framework to understand the emergence and spread of *de novo* mutations [23]. Viral dynamic modelling studies often employ the basic model of virus dynamics, the TIV model—a compartmental framework that tracks uninfected target cells (T), infected virus-producing cells (I), and free virus (V) (see refs. [24–28] for examples) or the TEIV model, an extension of the TIV model to include eclipse phase infected cells (E) that cannot produce virus particles (see refs. [29–36, 36–45] for examples). The TEIV model has emerged as a foundational model for acute viral infections to understand SARS-CoV-2 infection dynamics. The conceptual simplicity of the TEIV model and its analytical tractability make it widely applicable to many different families of pathogens; however, its assumptions warrant careful scrutiny when applied to mechanistically rich biological processes.

The standard TEIV approach models the eclipse and progeny-producing infected cells with the assumption of exponentially distributed lifetimes. These are biologically simplified assumptions, leading to virus-producing cells instantaneously capable of transitioning from a non-productive to a fully-productive state upon exiting the eclipse phase, and further means the eclipse phase can have an instantaneous (very short) duration. These are unrealistic assumptions given known intracellular delays [46]. Extensions of the TEIV models to apply non-exponentially distributed lifetimes are rare [36, 47–53]. Many biological processes involved in the viral host infection process occur through a series of sequential steps that result in non-exponentially distributed waiting times, often better approximated by gamma or Erlang, delay, or normal distributions [36], often necessitating more complex spatial modelling [54], or biphasic viral decay [55].

Investigating the mathematical oversimplification of an exponentially distributed eclipse phase has been of particular interest in the field of viral dynamic modelling, where for many pathogens a gamma-distributed eclipse duration has been deemed more reflective of the underlying biology [35, 45, 49, 51, 56, 57]. After a virus enters a cell, it undergoes a series of time-dependent steps such as uncoating, translation, genome replication, protein synthesis, assembly, and trafficking, before new progeny virions are ultimately produced [58]. The main biological motivation for investigating non-exponenital eclipse durations is to more realistically represent the timing variability of these intracellular processes. For example, Beauchemin *et al*. [59] found that using three or four eclipse stages is the most suitable approach to describe the duration of the eclipse phase during SHIV (Simian-Human Immunodeficiency Virus) infection *in vitro*. Liao *et al*. [28] investigated the number of stages in the eclipse phase needed to model Ebola and determined 13 stages would be needed. Furthermore, in the work detailed in refs.[38, 60, 61] on influenza virus infection, the number of stages included in the eclipse phase was set at 60, determined with an hourly unit duration, and in ref. [62] the most physiological value was determined to be 30. Further, in a recent study on SARS-CoV-2 infection, we found that at least three stages should be included [57]. These studies suggest that the eclipse phase is not a simple exponential but a biologically complex process whose accurate representation is essential for faithfully capturing viral dynamics across diverse pathogens.

It is clear from biophysical principles that the viral budding rate cannot instantly ramp from a state of zero budding virions to a constant budding rate. Biologically, this implies that newly activated infected cells begin shedding virus at their maximum rate immediately, without any intermediate build-up of the molecular machinery required for viral assembly and egress. For many pathogens, viral latency, shedding dynamics, and the influence of shedding dynamics on host immunity, are known to be important factors in successful host infection [63, 64]. For example, in the context of HIV infection models, mathematical frameworks have been advanced to determine optimal viral budding protocols that maximize viral fitness or better physiologically represent the viral shedding temporal dynamics [65–69]. Motivated to account for the fact that HIV virion production ramps up within the cytoplasm, Nelson *et al*. [69] investigate an age-structured target-cell-based model where the virion production and cell death rate are a function of the length of time of cell infection. They found that peak viral load levels, and time of peak, strongly depends on the progeny virion ramping protocol [69]. In the context of SARS-CoV-2, we are therefore interested in exploring the influence of virus ramp functions to account for the timescale and dynamics involved in reaching the maximally-budding virion state.

Whether viral dynamic work utilizing the standard TEIV model includes cell death during the eclipse stage, or virus loss during the infection of a target cell, often depends on the properties of the disease that is being modelled or what properties of the infection dynamics are of interest in the study. For example, if it is assumed that while the cell is being hijacked to produce virions, or has produced virions but none have yet egressed, that cytopathic affects are minimal, then cell death during the eclipse stage can likely be ignored [70]. For HIV, however, cell death during the eclipse stage is often considered because it is known that protein from infecting virus can be presented on MHC-1 molecules before the cell begins producing progeny virions, thus activating an immune response [71, 72]. For SARS-CoV-2, there is motivation to consider cell death during the eclipse phase [73], or budding ramp phases, because it has been shown that NK cells respond to early signs of inflamation at early stages of viral infection [74] and CD4+ T cells have been shown to be active within 48 hours of controlled viral inoculation [75]. Like the inclusion or omission of cell death during the eclipse stage, many models also include or omit virion loss, or absorption, from the infectious virion compartment. This term, when included, accounts for loss of free virions due to binding to target cells, or the rate at which virions irreversibly enter cells [76]. Studies have found that richer dynamics can occur when the virus loss term is included [29, 36, 57, 77–79].

Taken together, the body of work discussed above underscores the need for an extended TEIV model that can capture biologically realistic timing and loss mechanisms in the cell infection pathways. Such model choices can substantially influence the predictive power of within-host viral dynamics models. In this current work, we extend the TEIV framework by incorporating physiologically motivated ramp-up dynamics for viral budding, and fit our models to SARS-CoV-2 viral load trajectories. We investigate several mathematical schemas that describe how partially-productive infected cells gradually acquire the capacity to bud virus and reach a fully productive state Further, we consider viral load data drawn from predominantly hospitalized populations of individuals (severe infections), versus viral load data drawn from non-hospitalized individuals (mild infections). By fitting these extended models to SARS-CoV-2 viral load data, we evaluate their ability to better recapitulate observed viral kinetics compared to the standard TEIV model, and further, distinguish which parameter sets best represent severe (hospitalized) and mild (non-hospitalized) infection. Our results demonstrate that accounting for budding ramp up dynamics in viral budding improves model fidelity, offering a more accurate and mechanistically faithful representation of within-host viral dynamics. These refinements hold promise for improving the predictive modelling of infection progression and informing treatment strategies targeting different stages of the viral life cycle.

## 2 Methods

### 2.1 The TEIV Model

The standard TEIV model with *n* + 1 eclipse stages is given by the following equations:

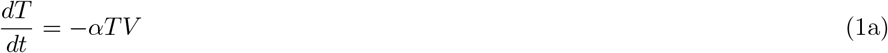

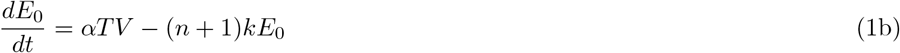

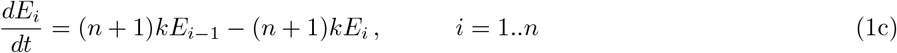

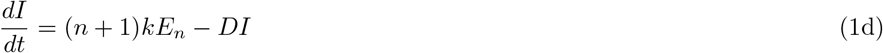

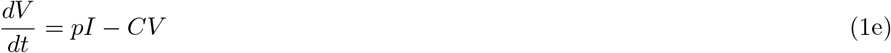

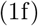

where *T* is the density of uninfected susceptible target cells, *E*_*i*_, *i* = 1..*n*, is the density of infected cells in the *i*^th^ eclipse stage, and *I* is the density of productively infected cells. Susceptible target cells are infected by a virus, *V*, at a constant rate *α*, and progress through the eclipse stages at rate *k*. The virus is produced by the productively infected cells, *I*, at a rate of p and is cleared at rate *C*. Productively infected cells die at rate *D* per cell. Model (1) assumes that an infected cell will transition from no virus budding/production to full virus budding immediately upon entering the productive stage, *I*. This assumption is unrealistic as a ramping up budding rate from zero to *p* should align with a cell’s intracellular processes [51, 59, 80, 81].

### 2.2 TEIV Model Extension to the TEAIV Model

In the current study, we expand on Model (1) to include an infected cell type *A*_*i*_, *i* = 1..*n* that has an increasing budding rate from zero in the *E* class to a maximal budding rate of *B* for infected cells, *I*. We refer to this model as the TEAIV model (see schematic in Figure 1). It also includes a Gamma distributed lifetime over the eclipse and ramp-up, *E* + *A*_*i*_ stages, and the virus loss term of infectious virus due to infection of a cell. The TEAIV model is given by

**Figure 1:**
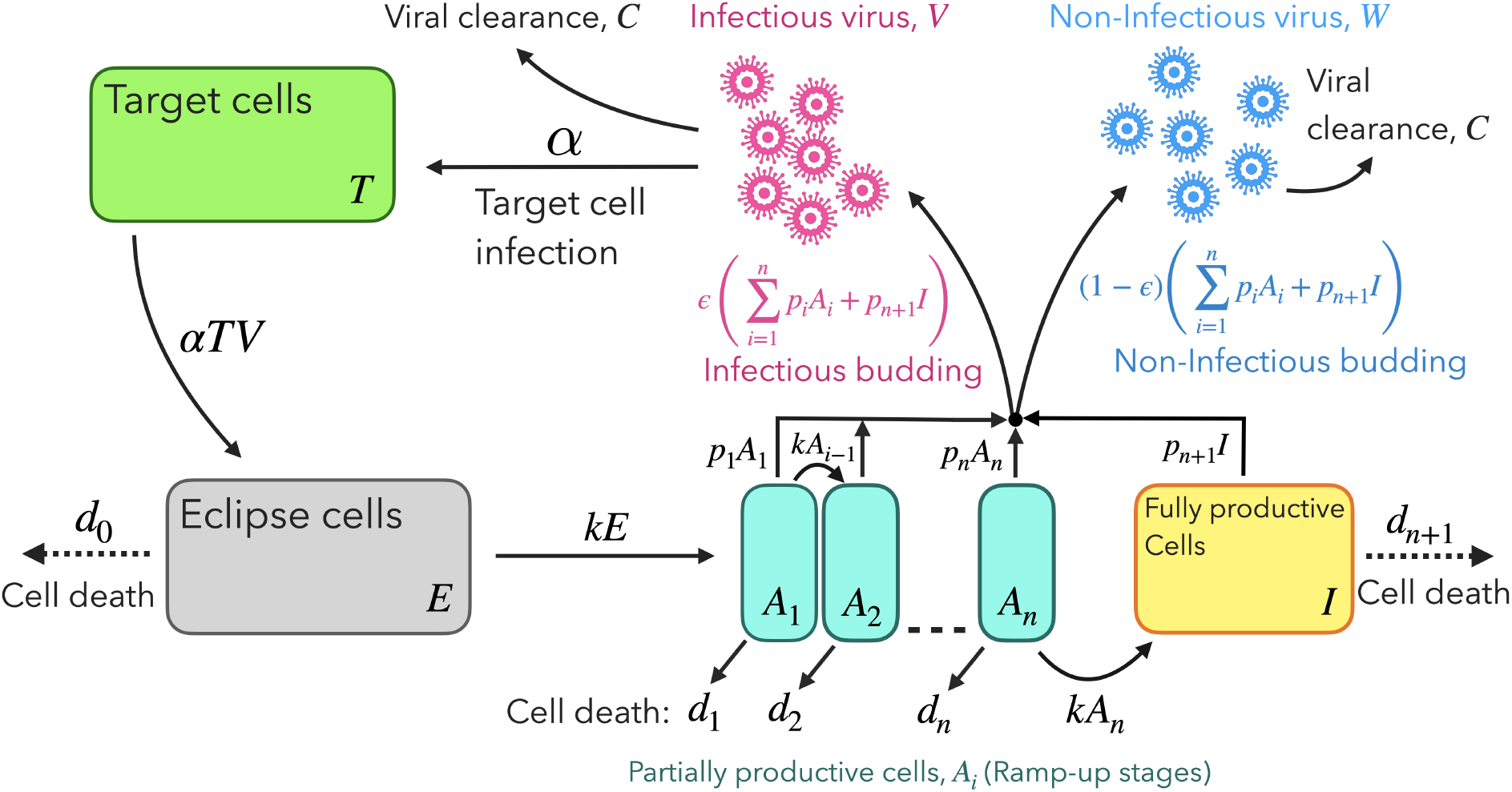
Schematic of the TEAIV model (Eq. (2)) displaying the dynamics of uninfected (T) and infected (E, A, I) target cells, and infectious (V) and non-infectious (W) virus. Partially productive cells, *A*_*i*_, ramp into maximally productive cells, *I*, via ramp function *p*_*i*_ *(Eq. (3)*). Cell death, *D, is dis*played by dashed arrow as it is both considered and not considered, see Section 2.4 for all model formulations. When cell death is considered, it is applied to eclipse (*E*), maximally productive (*I*) and partially productive (*A*_*i*_) cells.

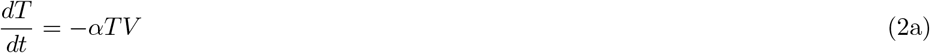

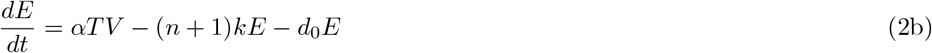

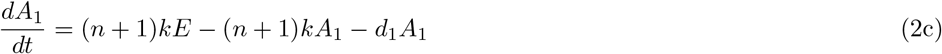

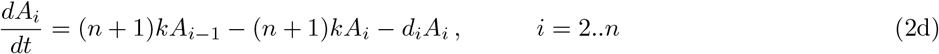

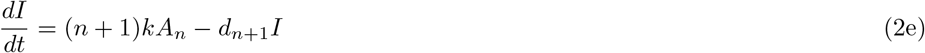

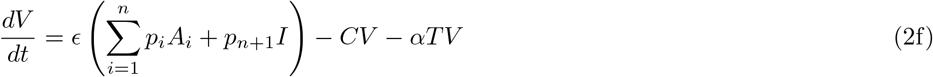

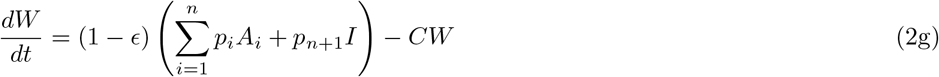

where

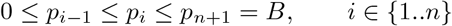

is the ramping up budding rate and

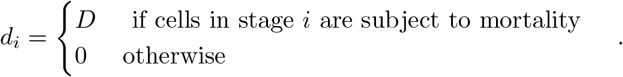

Model (2) describes the dynamics of uninfected (T) and infected (E, A, I) target cells, and infectious (V) and non-infectious (W) virus. Briefly, upon infection of an uninfected target cell *T*, at rate *α*, an infected cell *E* is produced whereby this cell cannot produce new virus particles. The newly infected cell then progresses to *A*_1_ and *A*_*i*_, *i* = 1..*n*, experiencing a ramping up budding rate of new virus particles (given by *p*_*i*_) after which it enters the fully productively infected cell class *I* with maximal virus budding rate *B*. Productively infected cells *A*_*i*_ (*i* = 1..*n*) and *I* can produce infectious *V* and non-infectious *W* virus where fraction *ϵ* are infectious. Virus particles are cleared at rate *C* and fully productively infected cells *I* can die at rate *d*_*n*+1_. Cell death, represented by *d*_0_ in the eclipse state, *E*, and *d*_*i*_ in budding stage *i, i* = 1..*n*, is considered in some fitting frameworks in this study (described below).

We note that virus production is influenced by the number of infected cells in each ramp-up stage *A*_*i*_, where *p*_*i*_ represents the fraction of the maximal budding rate *B* that cells in budding stage *i* produce. In our analysis, we present three distinct functions for *p*_*i*_^1^,

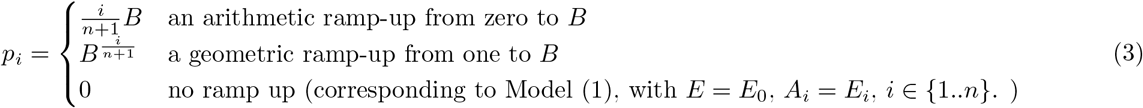

### 2.3 Virus Production

Since the duration of the eclipse plus ramp up stages is fixed, the effect of adding ramp up stages with arithmetic ramp up is a quicker ramp up (see Figure 2). Effectively, our arithmetic ramp up model is a stepwise increase in budding rate from 0 to *B*. Adding steps results in earlier budding, albeit at a low rate. In contrast, adding stages to our geometric ramp up model results in a slower initial ramp up, followed by a steeper ramp up in later stages reaching the full budding rate, *B*, sooner (see Figure 2, panel B). These observations imply that scenarios with more ramp up stages assume a quicker ramp up and an increase in the expected burst size (total virus released over the life of the infected cell) compared to scenarios with fewer ramp up stages. This phenomenon can be attributed to the introduction of additional intermediate steps in the process of viral replication and release. The eclipse phase is influenced by various intracellular processes, including viral nucleic acid and protein synthesis, viral assembly, maturation, budding, and successful release [82]. The presence of multiple eclipse stages leads to a more complex and intricate viral replication process, resulting in a quicker ramp up.

**Figure 2:**
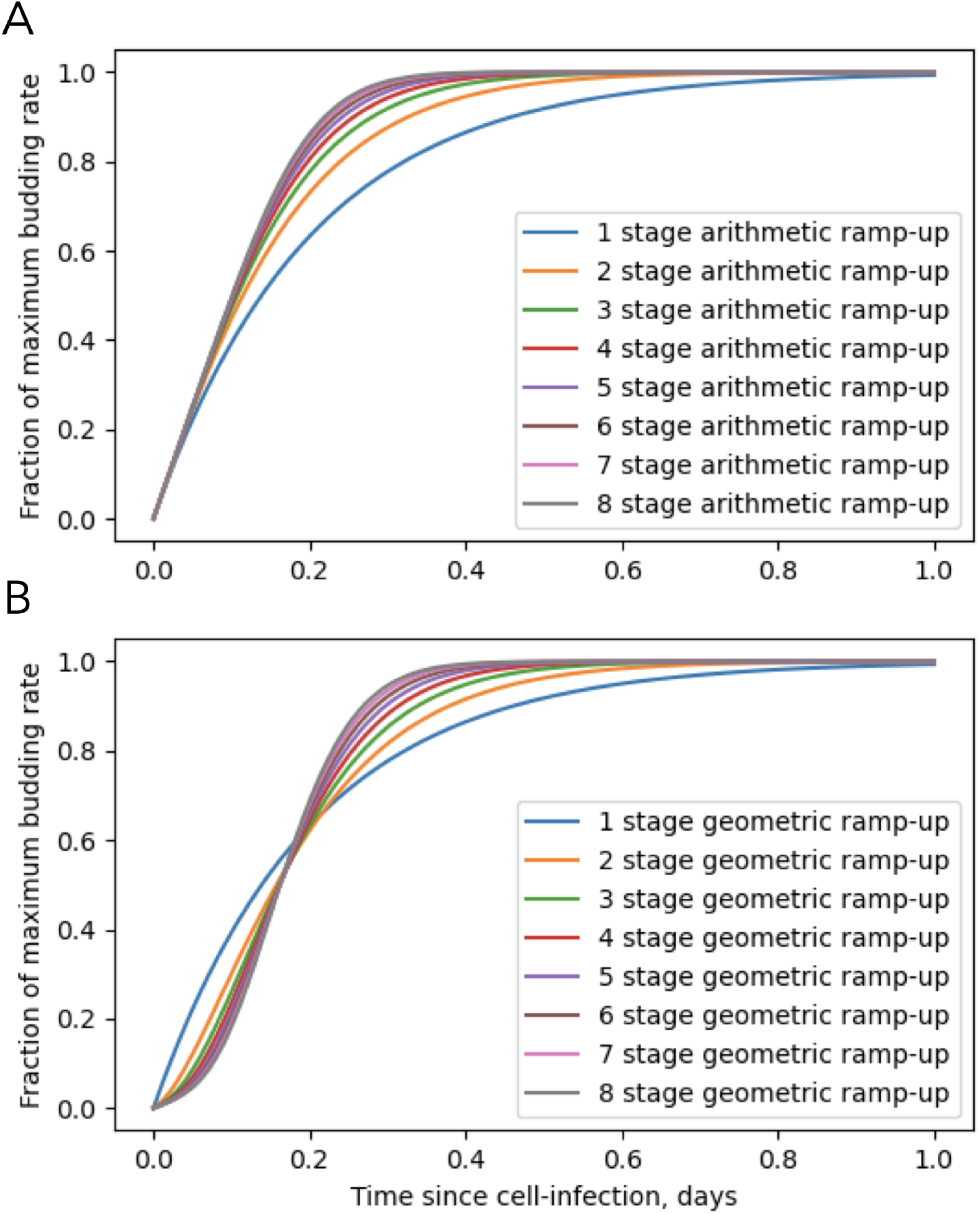
A) Fraction of maximum budding rate as a function of time since cell infection is shown for arithmetic ramp up models with 1 to 8 ramping stages, and B) Fraction of maximum budding rate for geometric ramp-up models for 8 stages, assessing the influence of the normalizing parameter *B*_0_. The curves are solutions of Model (2) with *α* = 0, *D* = 0, *B* = 1, and initial conditions of *E*(0) = 1 and all other states zero. These solutions effectively compute the budding rate of a single cell as it progresses through eclipse and infectious stages, barring cell death.

### 2.4 Model Fitting Frameworks

We fit the TEAIV and TEIV models to several SARS-CoV-2 viral load datasets to determine best fit models. We compare models that include or omit death rates in the eclipse state, *E*, and the ramp-up states, *A*_*i*_, *i* = 1..*n*. In total, we consider six model frameworks for the standard TEIV and newly presented TEAIV model:

1. No ramp-up <u>with</u> eclipse-state death (Model (2) with *p*_*i*_ = 0, *d*_0_ = *d*_*i*_ = *D, i* = 1..*n*).
2. No ramp-up <u>without</u> eclipse-state death (Model (2) with *p*_*i*_ = 0, *d*_0_ = *d*_*i*_ = 0, *i* = 1..*n*).
3. Arithmetic ramp-up <u>with</u> eclipse, *E* and ramp-up, *A*_*i*_ state death (Model (2) with 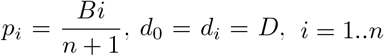).
4. Arithmetic ramp-up <u>without</u> eclipse, *E* and ramp-up, *A*_*i*_ state death (Model (2) with 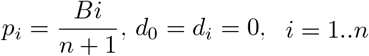).
5. Geometric ramp-up <u>with</u> eclipse, *E* and ramp-up, *A*_*i*_ state death (Model (2) with 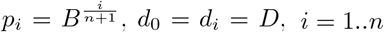.
6. Geometric ramp-up <u>without</u> eclipse, *E* and ramp-up, *A*_*i*_ state death (Model (2) with 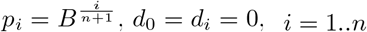).

For modelling frameworks 1-6 described above, we consider different forms of the Gamma distribution defining the lifetime of the cells in the eclipse state, *E*, and the ramp-up stages, *A*_*i*_, *i* = 1..*n*, and investigate *n* = 0..9, giving a total of 1 to 10 stages.

### 2.5 Initial Conditions and Parameter Values

Table 1 lists the initial conditions and parameter values for our study which align with previous studies of SARS-CoV-2 infection. The initial concentration of uninfected target cells *T* is assumed to be 1.3 *×* 10^5^ cells mL^*−*1^ [35, 57, 83, 84]. We assume that there is one productively infected cell present in the upper respiratory tract at time zero, specifically at a concentration of one cell per 30 milliliters (mL) of fluid [57], and assume that there are 10^*−*7^ newly infected cells *E* [57]. There are zero virus particles and zero ramp-up stage, *A*_*i*_, *i* = 1..*n* cells at time zero.

**Table 1:** Model parameters and initial conditions. NA denotes unitless. * denotes informed priors for model fitting with random effects turned on.

| Parameter | Definitions | Value | Unit | Comments |
| --- | --- | --- | --- | --- |
| $\alpha$ | Rate of attachment per target cell | $4.93 \times 10^{-5*}$ | $\frac{mL}{day.copies}$ | Fixed across population[83], random effects estimated |
| k | Reciprocal of the total duration the eclipse, $E$, and ramp-up, $A_i$, $i = 1..n$ stages | 5 | $\frac{1}{day}$ | Fixed, [85] |
| D | Loss rate of infected cells | 1* | $\frac{1}{day}$ | Fitted |
| $\epsilon$ | Portion of active virus particles made by budding cells | 0.001 | NA | Fixed, [83, 84] |
| B | Maximum rate of virion release from infected cells | 8000* | $\frac{copies}{day.cell}$ | Fitted, [83] |
| $p_i$ | Rate of virion release in stage $i$ | Eq. (3) | NA | Determined by the Zero Budding, Arithmetic Ramp-Up or Geometric Ramp-Up functions |
| $C$ | Rate of virion clearance | 10 | $\frac{1}{day}$ | Fixed, [35, 57] |
| $t_{inf}$ | Time of initial infection | 1* | day | Fitted |
| $T_0$ | Susceptible target cell initial condition | $1.33 \times 10^5$ | $\frac{cells}{mL}$ | Fixed, [57, 83] |
| $E_0$ | Newly infected cell initial condition | $10^{-7}$ | $\frac{cells}{mL}$ | Fixed, [57] |
| $A_{i0}$ | $i$th eclipse state initial condition | 0 | $\frac{cells}{mL}$ | Fixed, [57, 83] |
| $I_0$ | Budding cells initial condition | $\frac{1}{30}$ | $\frac{cells}{mL}$ | Fixed, [57, 83] |
| $V_0$ | Infectious virus initial condition | 0 | $\frac{cells}{mL}$ | Fixed, [57] |
| $W_0$ | Non-infectious virus initial condition | 0 | $\frac{cells}{mL}$ | Fixed, [57] |

Model (2) includes at least seven and at most seventeen parameters, corresponding to the model with one or ten ramp-up stages, respectively. Some parameter values are informed by the literature, while others are determined from model fitting to clinical cohort data. Table 1 identifies which parameters are fit to data versus fixed to SARS-CoV-2 infection literature-informed values. Informed priors for the model fitting are identified by an asterisk (*). Note that the fitted value of *t*_*inf*_ is constrained so that infection occurs before time zero (the data of first patient sample) and it is bounded between 0 and 14 days, similar to [57, 83], if the pre-peak viral load data is not present.

### 2.6 Clinical data

We fit our models to previously published hospitalized [83, 86, 87] and non-hospitalized [88] viral load patient data. In all datasets, viral load counts are in log_10_ units/mL. The Néant [83] dataset contains information from 655 patients who were hospitalized, with a median age of 60. Nasal swabs were used to measure the viral load. The viral load measurements in Jones et al. [86] come from 4344 people with a median age of 52, of whom 3475 were hospitalized. Measurements of viral load were obtained using sputum swabs. The Goyal et al. [87] study includes 25 people whose viral load time courses were measured using throat and nasopharyngeal swabs. Finally, in the non-hospitalized dataset [88], data was collected from 60 individuals for up to 14 days. Measurements of viral load were collected using nasal swabs and saliva samples. The median age of the cohort is 28 years; range, 19–73 years. We note in all datasets no individuals reported receiving the SARS-CoV-2 vaccine prior to participation, and all datasets come from fairly early on in the COVID-19 pandemic where the wildtype and B117 variants likely dominated.

### 2.7 Fit procedures and goodness of fit criteria

We fit our mathematical models to the clinical data using Monolix [89]. Monolix employs a maximum likelihood estimation method, finding the values of the parameters that maximize the likelihood of the observed data, given the model. In our study, every fit begins with similar priors and every model has the same number of fitted parameters. To assess model goodness of fit to the data, we employ Monolix’s error model “combind1” (combined error model) and we assume that the residuals follow a normal distribution. Following the methodology outlined in [83], our model evaluation process takes into account both the Bayesian information criterion (BIC) value and the relative standard error (RSE). The process of selecting the best-fit model is guided by several additional factors, qualitative aspects of the goodness-of-fit plots, accurate parameter estimates as informed by the literature, and physiological plausibility of parameter values.

The BIC is a measure of model fit that penalizes the number of parameters in a model to reduce the risk of overfitting. Lower BIC values indicate better model fits [90]. The candidate model with the highest preference is the one with the lowest BIC (*BIC*_*min*_) among all of the candidate models. When comparing BIC values, a ΔBIC > 2 is considered to be significant [91].

Relative Standard Error (RSE) provides a measure of the relative uncertainty or variability associated with the estimated parameters in a statistical model. The RSE is calculated as the ratio of the standard error of a parameter estimate to the estimate itself. It is often used to assess the precision or reliability of parameter estimates. A lower RSE indicates a smaller relative uncertainty or variability, suggesting a more precise estimate. The RSE is a useful metric for evaluating the quality of parameter estimates. According to the Central Bureau of Statistics (2018) [92], an estimate is considered accurate for publication with an RSE less than or equal to 25%.

## 3 Results

### 3.1 Hospitalized Data

#### Model Fitting

We fit the TEIV and TEAIV models with and without death rate *D* in the *E* and *A*_*i*_ classes, to datasets from refs.[83, 86, 87], as described in the methods. For each model, fitting to all datasets is conducted simultaneously within Monolix. The resulting values for the relevant viral kinetic parameters are listed in Tables S1 - S3. We note that all model fits employ an identical count of parameters and share comparable initial values, as detailed in Table 1. The resulting model fits with *n* = 7 with cell death are shown in Figure 3 alongside the clinical data. The values of the model fit parameters are listed in Table S4.

**Figure 3:**
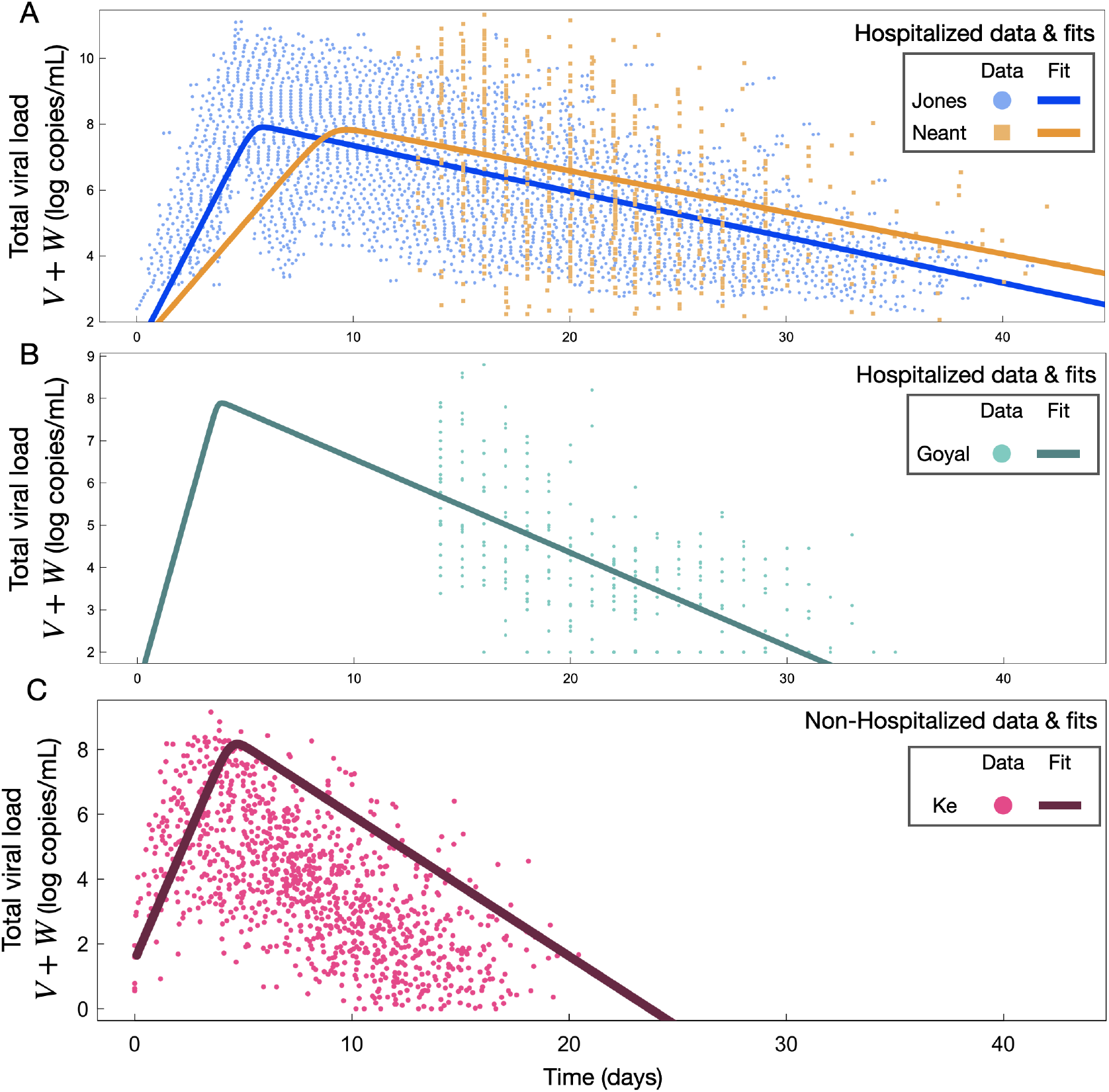
Model fit examples. The total viral load, Eqs. (2)f+ (2)g (*V* + *W* ), are shown for lowest BIC TEAIV model fits for A-B) the hospitalized (severe-case) data corresponding to the TEAIV model with arithmetic ramp up, *n* = 7, with cell death for the Neant [83] and Jones [86] (A) datasets and Goyal [87] (B) datasets; (C) the non-hospitalized (mild-case, Ke [88]) data corresponding to arithmetic ramp up, *n* = 9, with cell death. (A-C) are depicted on the same x-axis scale for ease of comparison.

#### Model Selection

Table 2 displays the BIC (Bayesian Information Criterion) value for the model fits to the hospitalized datasets and the relative standard errors on the fitted parameters. We observe that models including the arithmetic ramp-up budding rate provide the minimum BIC value for all models. Considering the model with arithmetic ramp-up only, we find that the TEAIV model with death rate *D* and seven total eclipse stages has the smallest BIC value (Table 2, bold text). As ΔBIC > 2 is deemed significant [91], for arithmetic ramp up we determine that all models with more than two ramping stages are accepted as best model fit. We note that all BIC values for the TEIV models and the TEAIV models with geometric ramp up have ΔBIC *>* 2 compared to the minimum value for arithmetic ramp up. We also note that all but one TEAIV model with arithmetic ramp-up have a ΔBIC < 2 compared to the minimum BIC. Finally, we observe that the TEIV and TEAIV models with geometric ramp-up all have BIC values with ΔBIC values greater than two compared to their TEAIV artimetic ramp-up model counterparts (comparing results across columns, Table 2). Table 2 lists the RSE of all model fits for fitted parameters *D, B* and *t*_*inf*_ with the minimal values highlighted that also correspond to the minimal arithmetic ramp-up BIC models. The max/min RSE on *D* and *B* are found to be 28.5/17.3% and 122/22.9%, respectively, across all arithmetic model model fits, and the max/min RSE on the initial time of infection is found to be 32.7/6.12%.

**Table 2:**
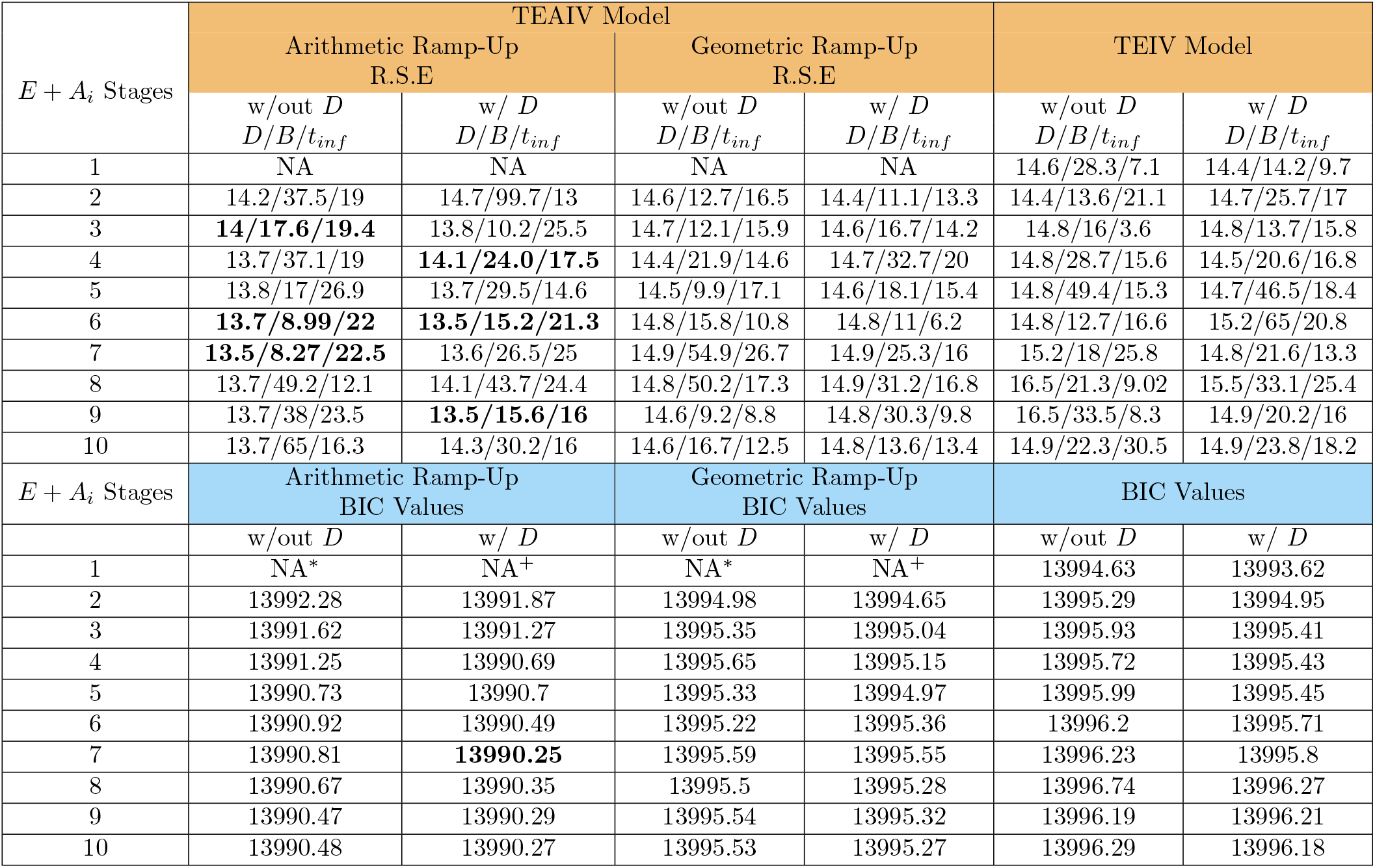
Top (orange header): Relative Standard Errors (RSE) for Estimated Parameters from severe infections (hospitalized individuals). The RSE value is determined for the population estimate of each parameter. Each row represents a Monolix run with change in eclipse stage number, inclusion of death rate D in the *E* and *A* classes, and ramp up function. NA^*∗*^ and NA^+^ denote that these models are the sames as the TEIV model without (w/out) and with (w/) *D* in the *E* class, respectively. Bottom (blue header): BIC values for severe infection (hospitalized individuals) } . Each row represents a Monolix run with change in eclipse stage number, inclusion of death rate D in the *E* and *A* classes, and ramp up function. NA^*∗*^ and NA^+^ denote that these models are the sames as the TEIV model without (w/out) and with (w/) *D* in the *E* class, respectively

Given the BIC values (Table 2), we conclude that the Arithmetic Ramp-up models offer the best balance between goodness of fit and model complexity, making them favorable choices for explaining the observed data for the hospitalized dataset. Given the RSE values (Table 2), we find that arithmetic ramp-up model 3 without *D*, model 4 with *D*, model 6 with or without *D*, model 7 without *D*, and model 9 with *D* have RSE values lower than 25% (Table 2, bold text). We thus conclude that these models provide the best models to explain the hospitalized clinical cohort data.

### 3.2 Non-hospitalized Data

#### Model Fitting

The TEIV and TEAIV models are fit to the non-hospitalized clinical data. A example of the best-ft model (described below) is shown in Figure 3C. The values of the model fit parameters are listed in Table S4.

#### Model Selection

Similar to that conducted for the hospitalized datasets, we proceed to select models that best represent the non-hospitalized data. Table 3 lists the BIC and RSE values determined from our model fitting. Similar to the hospitalized data fits, we also find that the arithmetic ramp-up TEAIV model provides the most minimal BIC, with ΔBIC*<* 2 compared to geometric ramp-up and the TEIV model. The most minimal BIC model is determined to be the arithmetic ramp-up model with 8 or 9 ramp stages, however, we note all models with ramp stages greater than 1 are equivalent via BIC comparison. Across all arithmetic ramp-up models the max/min RSE on *D* and *B* are 26.4/17.5% and 122/14.2%, respectively. The max/min RSE on the time of infection fit for the arithmetic models is 32.7/7.88%. We highlight in bold the minimal RSE parameter combinations, including the 9 ramping stages fit which is determined to be one of the minimal BIC models. Overall, we find that arithmetic ramp-up with 9 stages, with or without D, to be the best fit model for the non-hospitalized dataset, the model fit with D is shown in Figure 3B.

**Table 3:**
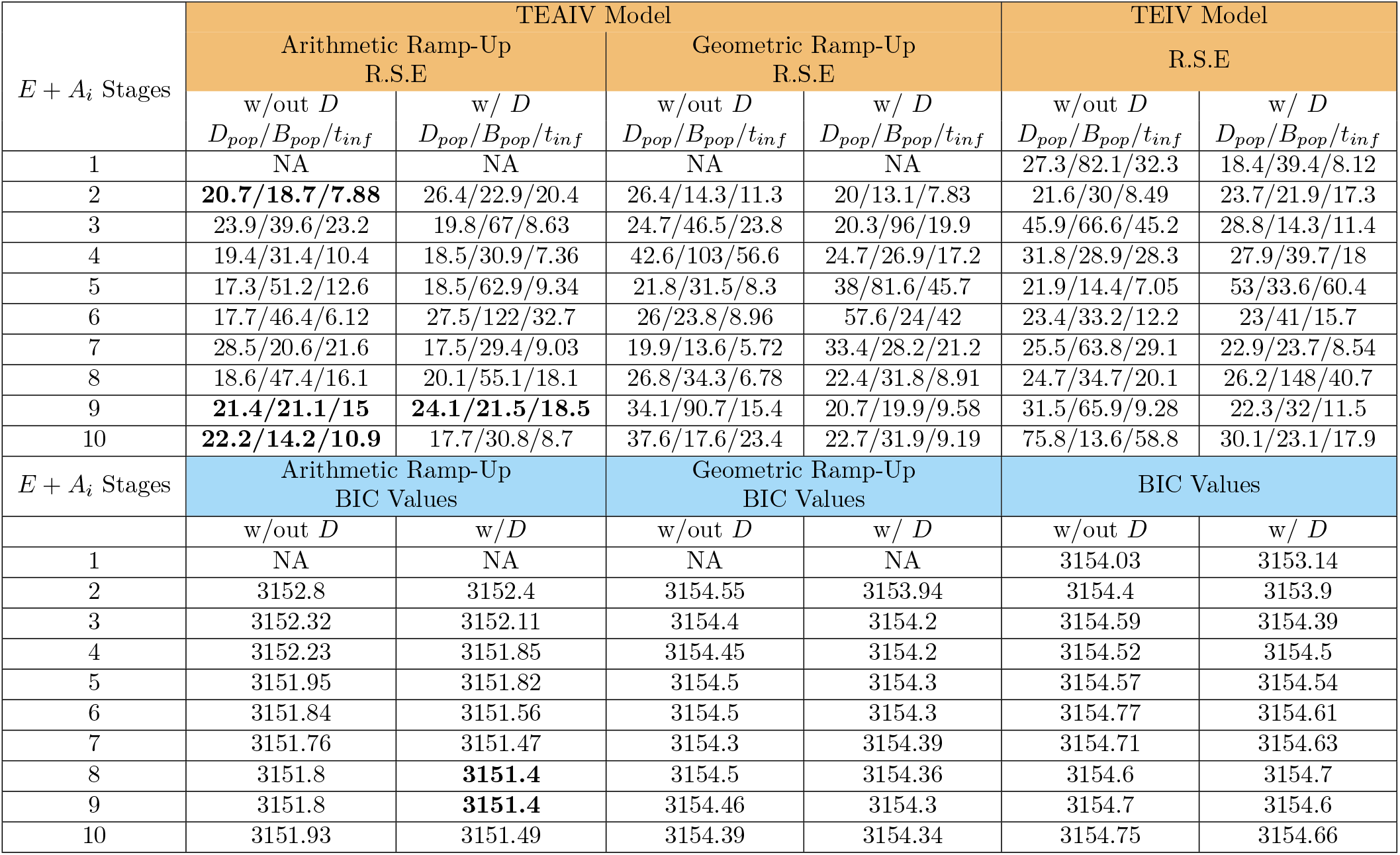
Top (orange header): Relative Standard Errors (RSE) for Estimated Parameters from mild infections (non-hospitalized individuals) in Monolix Models with Added Eclipse Stage, Death Rate (D), or Ramp Up Functions. *B*_*pop*_ : population estimation of virion budding rate from infected cells, *D*_*pop*_: population estimation of death rate, *t*_*inf*_ : population estimation of the time of infection. Bottom (blue header): BIC values for mild infection (non-hospitalized individuals). TEIV and TEAIV model outcomes for all model fitting frameworks described in Section 2.4 as the number of eclipse stages are varied from 1 (exponential) to 10 (Erlang). The lowest BIC highlighted, however, note that ΔBIC within 2 are equivalent.

### 3.3 TEAIV Model Reproduction Number

In Supplemental Section 1.1 we provide a derivation of ℛ _0_ for the general TEAIV model with cell death as a function of ramping state, arbitrary ramping state transition times, and arbitrary ramp function *p*_*i*_. For the special TEAIV model cases considered for fitting in this manuscript, we find ℛ _0_ for Model (2) with cell death applied to eclipse and ramping to be,

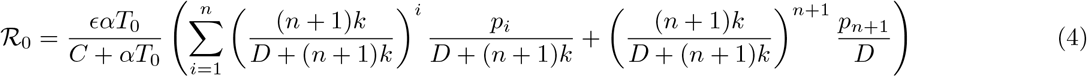

and we determine ℛ_0_ for Model (2) without cell death in the eclipse and ramp-up stages to be,

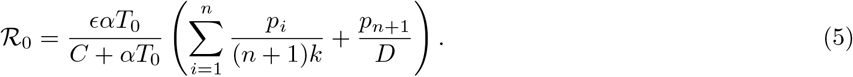

The interpretation of the TEAIV model ℛ_0_ is similar to that of previous models: the probability of effective viral transmission per virus particle is multiplied by the total viral production, providing the average number of secondaryc infections generated by one primary infection in a fully susceptible population. Here, we breakdown the specific contributions to the TEAIV model ℛ_0_ considered in this paper. The prefactor, *ϵαT*_0_/(*C* +*αT*_0_), is interpreted as the probability that a newly produced effective (*ϵ*) virus particle will successfully infect a target cell rather than being cleared; this term balances the rate of effective infection against the total rate of virus removal (both by clearance and by successful infection). For both Eq.’s (4) and (5) the term in brackets following the prefactor represents the burst size: the total viral production capacity per infected cell, incorporating the viral load contribution of each partially infected cellular compartments (*A*_*i*_) via ramping function *p*_*i*_ as well as the contribution of the fully infected state (final term in brackets), via the contribution *p*_*n*+1_. Here, *p*_*i*_ is an arbitrary ramping function. In the case where cell death is possible for all partially productive states *A*_*i*_ (Eq. (4)), the term ((*n* + 1)*k/D*(*n* + 1)*k*)^*i*^ represents the probability that an initially infected cell successfully reaches compartment *A*_*i*_ (passing through E and *A*_1_ to *A*_*i−*1_) without dying, while *p*_*i*_*/*(*D* + (*n* + 1)*k*) represents the infectious viral load yield from the *i*^th^ partially infected cell state, *A*_*i*_, before either progressing to the (*i* + 1)^st^ state or dying. The final term in Eq. (4) represents the effective virion contribution from the fully productive cell, *I*: ((*n* + 1)*k*/(*D* + (*n* + 1)*k*))^*n*+1^ represents the probability that an initially infected cell successfully reaches the fully productive state, passing through *E* and *n A* compartments (*n* + 1 successful transitions), and *p*_*n*+1_/*D* is the virion production from the fully productive cell multiplied by 1/*D*, the mean time spent in the fully productive state. Since *I* is the terminally infected, or fully productive, compartment, its cells only decay (not progress to another compartment), so death rate *D* is the only contribution to the denominator for the fully infected virion production term. In the case where cell death is *not* possible for all partially productive states *A*_*i*_ ℛ _0_ is given by (Eq. (5)). Here, the terms contributing to the burst size have the same meaning as in Eq. (4) but are simplified. The probability that a cell successfully reaches compartment *A*_*i*_ is 1.0, and *p*_*i*_/(*n* + 1)*k* is the infectious viral load yield from the *A*_*i*_ partially infected cell state before progressing to state *A*_*i*+1_. Where cell death is applied only to the fully infected state, which is reached with probability 1.0, *p*_*n*+1_/*D* is the virion production rate scaled by death rate, *D*.

In the main text of this paper we consider *p*_*i*_ given by arithmetic and geometric functions (Eq. (3)) and with constant (or zero) cell death applied to the ramping states. In the supplementary material we analytically consider the most general TEAIV model with arbitrary *p*_*i*_, arbitrary cell death for each compartment, and arbitrary transition rates between partially-productive cell states (Eq. (S1)) leading to a reproduction number for the general case, given by Eq. (S5). From the general model we also analytically consider a hill function saturating ramping process, where in the continuum limit the compartmental TEAIV model with *n* compartments transforms to a PDE with drift (Section S1.2). We save for future work model fitting and BIC minimization to these general TEAIV models.

Scatter plots of ℛ _0_ as a function of the number of ramping stages are shown in Figure 4. Similar to previous work fitting TEIV models to hospitalized and non-hospitalized data [57] we find that for the non-hospitalized dataset, which lacks information on peak magnitude, peak placement, and pre-peak initial data (see Figure 3B), the resulting ℛ _0_ show significant variance as a function of the number of ramping states, *n*, for all model considerations (Figure 4A). Conversely, ℛ _0_ values determined from model fits where the full viral load trajectories are present with highly resolved peaks (see Figure 3A) are found to collapse to very similar values across all TEAIV model considerations (Figure 4B). The corresponding burst size calculations for all considered models is provided in Figure S1.

**Figure 4:**
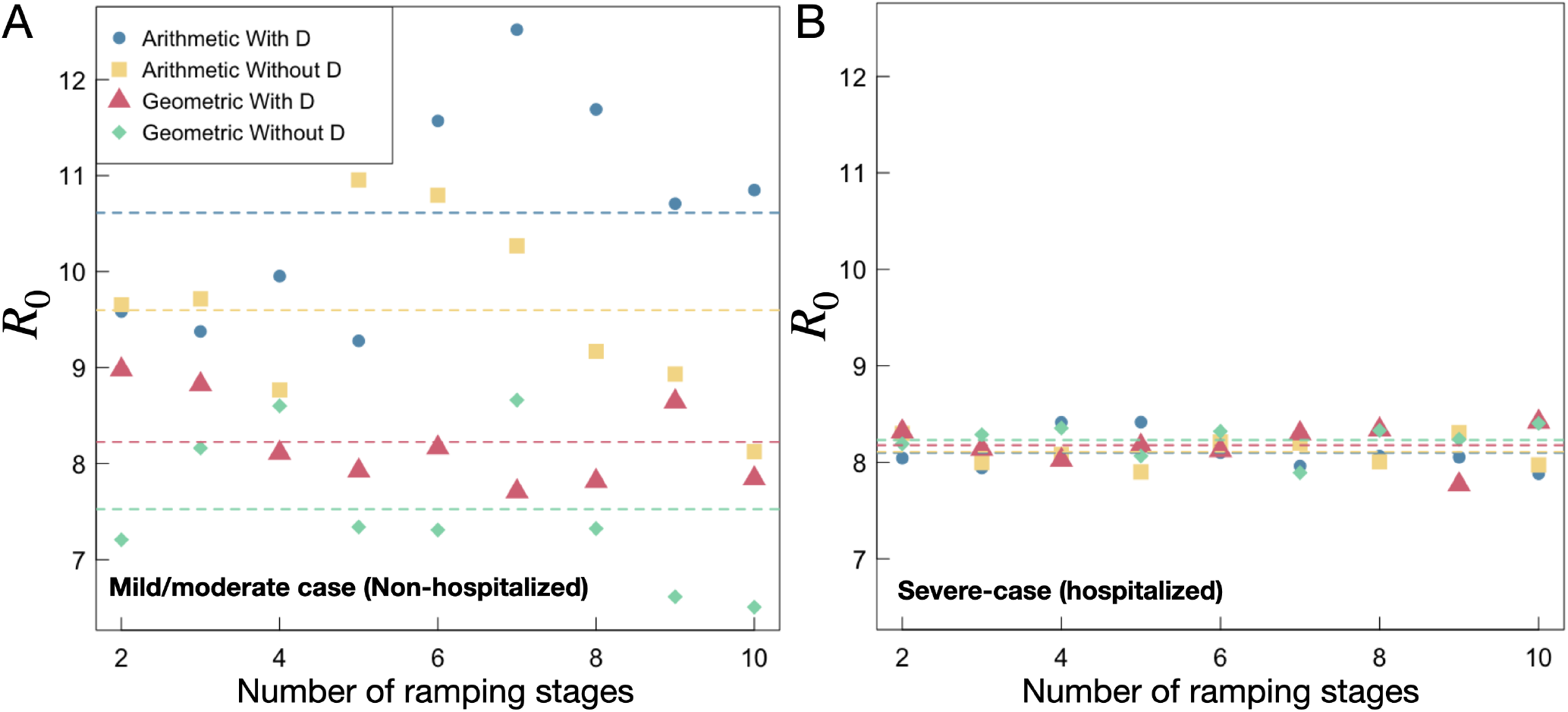
ℛ_0_ **numerical analyses computed from Eq.’s** (4) **and** (5). ℛ_0_ as a function of number of ramping stages for (A) the mild/moderate case (non-hospitalized) and (B) the severe case (hospitalized) TEAIV model fits.

## 4 Discussion

In this work we extended the TEIV model to include viral budding ramp up in a new model framework, the TEAIV model, where instead of *n* + 1 eclipse compartments we investigate one eclipse compartment plus *n* budding cell compartments, *A*_*i*_, that each contribute to the production of progeny virions. We investigated two general mathematical approaches to describe virion budding ramp up, arithmetic and geometric (Eq. (3)), and fit these models to four previously published SARS-CoV-2 viral load datasets. We also investigated the influence of varying the number of eclipse stages from 1 to 10 in the TEIV model, to compare to 1 to 10 virion-producing ramp stages, to test if there is a clear minimal BIC across all ramp up and eclipse stage model frameworks in TEIV and TEAIV formulations. For each of the just-described formulations, we further explore whether including cell death in the eclipse, or ramping, compartments influences the model fits as determined through the BIC and relative standard error. Finally, the viral load data is distinguished between predominantly drawn from predominantly hospitalized individuals (severe infection) versus non-hospitalized individuals (mild infection).

### The TEIV model

Following the successful attachment and entry of an infectious virion into its target cell, progeny virus is usually not observed for a delayed period, known as the eclipse phase (we note that this definition differs from the ‘period of infection to time of first possible detection’ [93]). Physiologically, the eclipse phase generally describes the myriad of viral hijacking mechanisms that must occur prior to the production of viral progeny (e.g., transcription, translation, genome replication, assembly, and trafficking) [46]. Mathematically, the delay before production of progeny virus following successful virion entry was originally modelled by considering two separate populations of infected target cells: *eclipsed, E* (infected but not yet viral-budding) lasting for a mean period 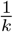, and productively infected, *I* (producing viral progeny at some rate, *p*). The TEIV model, consisting of a single eclipse compartment, leads to an exponentially distributed eclipse duration. This is an issue, originally raised in the modelling of SHIV [50], whereby an exponentially distributed eclipse duration mathematically means cells are capable of instantaneously budding virus upon viral entry, a physiological impossibility. Thus, much work on accurate modelling of the eclipse duration and shape has been advanced for a number of viral pathogens such as, but are not limited to, SARS-CoV-2 [57, 94], Zika [70], SHIV [50, 56], RSV [95], and influenza [38].

In the current study, we advance eclipse-duration models applied to SARS-CoV-2 by investigating the TEIV model with up to 10 eclipse stages, both with and without cell death rate, *D*, incorporated in the eclipse compartments, and compare to TEAIV models that allow for up to 10 budding ramping compartments, from zero to a maximal budding rate of virus particles per time unit. We then look to see which specific model minimizes the BIC and leads to physiologically defendable parameter values. Models *with* and *without* cell death, *D*, are examined because it is not well known whether immune-mediated clearance occurs during the eclipse phase of infection, particularly during early infection times where NK cells have been shown to respond to early signs of inflammation at early stages of viral infection [74] and CD4+ T cells have been shown to be active within 48 hours of controlled viral inoculation [75]. Incorporating a death rate for eclipse-phase cells in mathematical models may allow for a more accurate representation of the immunological processes involved in viral infections; it acknowledges that not all infected cells survive to become productively infected and that immune-mediated clearance can occur at various stages of infection. Our hypothesis was that this addition may improve the model’s ability to fit empirical data and enhance predictive power.

For the TEIV model fit to severe-infection viral load data, we find that both including or excluding cell death in the eclipse compartment, tested across 1 to 10 eclipse stages, leads to nearly-indistinguishable model performance with all ΔBICs falling within 2 across all TEIV model fits (Table 2, right-most two columns). A very slight preference, and minimal BIC, is found for the single eclipse stage model; however, we caution that when rounded, the BICs are equivalent. Similarly, for the TEIV model fit to mild-infection viral load data all ΔBICs fall within 2 (Table 3, right-most two columns). Therefore, incorporating cell death during the eclipse phase, even across increasingly fine-grained stage structures, does not significantly improve model fit (as assessed via BIC), for the TEIV model.

### The TEAIV Model

From a physiological standpoint, the transition of an infected cell from non-productive state to fully virus-producing is not instantaneous. After, or during, the eclipse phase, intracellular processes such as viral gene expression, protein synthesis, genome replication, assembly, and virion trafficking require time before a cell reaches its maximal virion output capacity. Therefore, the common modelling assumption that infected cells immediately begin shedding virus at a constant rate oversimplifies the true biological dynamics. This is particularly important when interpreting viral load data, which are typically measured from blood or saliva samples taken over time in a cohort of individuals. Aggregate viral load measurements reflect the cumulative behaviour of large populations of infected cells, where gradual ramp-up in viral budding across many cells may manifest as smoother time-dependent changes in observed viral load. From a mathematical perspective, our models use ordinary differential equations to track concentrations of infected cells and free virus over time, representing the average behaviour of cell populations rather than individual cellular trajectories. In this framework, we implemented arithmetic and geometric viral budding ramp functions to model a gradual increase in the per-cell virus production rate after the eclipse phase (Model (2)). The TEAIV model provides a tractable way to smooth the onset of viral output over non-instantaneous timescales, avoiding biologically implausible discontinuities in viral shedding. Our hypothesis is that incorporating gradual ramp-up dynamics can improve the model’s ability to capture infection kinetics.

To assess the impact of our model choices, we examined geometric and arithmetic (Eq. (3)) ramping behaviour. Like in the TEIV model study, we conducted the model fitting both with and without cell death incorporated in the budding ramp up compartments. Here, we determined that an arithmetic ramp-up framework leads to the most minimal BIC, specifically if the *E* + *A*_*i*_ stages exceeds 3 (Table 2), regardless of presence of cell death in the viral budding ramp compartments, which are determined to be equivalent models via BIC. We note that arithmetic ramp up also minimizes the BIC when comparing to all the TEIV formulations considered (Tables 2 and 3).

When the TEAIV model is fit to severe-infection viral load data, we find that the BIC minimizes beyond 3 ramping stages for arithmetic ramp up (Table 2). For the geometic ramp up, we find that all fits remain within ΔBIC of 2.0. Overall, for model formulations fit to hospitalized data, the TEAIV model with arithmetic ramp-up and ≥ 3 ramping stages leads to the most minimal BIC. We find similar outcomes for the non-hospitalized data, where the best-fit model with most minimal BIC are the TEAIV arithmetic ramp-up models with more than 3 ramp-up stages.

Altogether, given that models with and without ramping-associated cell death are statistically indistinguishable (via BIC), simpler models may be preferred for practical use when modelling SARS-CoV-2 dynamics. In light of the best explanatory results for *i* > 3 and the concerns of identifiability with increasing *n* and thus model complexity, we remark that the choice of the number of eclipse or ramping stages is one of model convenience to introduce budding ramp-up. Additionally, biologically budding may not occur in explicitly discrete stages but ramp-up continuously as the infected cell matures to a fully-infected state. As such, a large *n* continuum limit may be a more appropriate tool to analyze this process. This would replace discrete budding stages *A*_*i*_ and budding rates *B*_*i*_ with cell densities *A*(*x*) and rates *B*(*x*) with a continuous infection maturation rate *x*. This has the advantage of the biological complexity of multi-stage ramping but the mathematical simplicity of reducing a series of *n* compartmental differential equations for *A*_*i*_ to a single partial differential equation for *A*(*x*). This may have further benefits in addressing parameter identifiability. The details of a continuous limit for the model following a Hill function ramp-up are detailed in Section S1.2 of the Supplementary Material.

### Variability in Data-Fitted Parameter Values by Data-Set

All fitted parameter values across all datasets for the TEIV and arithmetic/geometric TEAIV models are given in supplementary Tables S1 to S4 with population-level parameters provided in Table S4.

Similar to previous in-host studies on SARS-CoV-2 [57] with data from the studies of Neant [83], Goyal [87], and Jones [86], we observe differences in best-fit parameter values between the datasets (Tables S1 to S3, column *D*). We however note that within each dataset, irrespective of model formulation (TEIV or (TEAIV), budding ramp-up structure, and inclusion or disclusion of *D*, the respective fitted values for cell death, *D*, are relatively consistent – *D* is determined to be approximately 0.29, 0.51, 0.32, across all *E* and *A* stages, for Neant, Goyal, and Jones, respectively. Considering, the fitted value for *α*, we also see some consistency within datasets across all models. The fitted budding rate, *B*, however, seems to be more sensitive to model and ramp-up structure, and the inclusion of *D*. The largest shift in budding rate is found by comparing the TEAIV framework with versus without *D* incorporated in the eclipse/budding stage; specifically, the budding rate *B* is highest (exceeding 13000) when employing arithmetic ramp up with *D*, but only for the Neant and Goyal datasets where viral load peakedness is not highly resolved (Figure 3), otherwise, *B* is approximately 8000 similar to the other model fits described above. These data-source dependent results may be related to structurally-identifiability effects from the various models, or, as the datasets have variable resolution of viral dynamic kinetics, these results may be due to variability in the data resolution of the pre-peak viralload growth phase, previously shown to influence practical identifiability of TEIV-based model parameters [57, 96]. Differences in the temporal resolution and quality of viral load measurements across experimental datasets can lead to variability in how tightly model parameters are constrained, affecting identifiability and the robustness of inferred dynamics.

In examining the non-hospitalized cohort (Figure 3C), our TEAIV framework was unable to reproduce the sharp post-peak decline in viral load that is consistently observed in this dataset. One plausible explanation, supported by recent modelling of the same dataset, is that the presence of a refractory cell compartment is needed to capture the dynamics, whereby interferon signalling induces a transient antiviral state in susceptible cells, preventing their infection while preserving the pool of target cells [88]. In the refractory cell model, binding of interferons converts a fraction of susceptible cells into refractory cells that are temporarily protected, such that the susceptible cell population is not rapidly exhausted. Using a model with refractory cells, *Ke et al*. [88] reported an individual rapid infected cell death rate of ∼2 −3 per day with a mean population estimate of 2.45 per day. This is significantly faster than the value found from our TEAIV models (0.75 per day, Table S4). We will consider an extension of the TEAIV model to incorporate the effects of refractory cells in future work.

Future work will consider other ramp-up models that are influenced by the timescales of the immune response or explicitly couple virus production to immune dynamics. For example, future efforts could incorporate viral budding rates that depend on cell/tissue damage accumulation, local cytokine concentrations, and interferon signalling thresholds [97, 98]. Models could also include spatial effects that allow for the explicit modelling of varying cell characteristics, viral budding rates, and immune responses in tissues of the upper and lower respiratory tracts (see [1] for some examples). Over all these extensions, a variety of model frameworks can be considered, including delay differential equations, dynamic threshold functions, and spatially structured compartments. Of immediate interest is an extension of our model to a partial differential equations framework that includes a continuous budding rate, and a study of the implications that this may have on parameter estimation and identifiability.

In the current work, we limited our study to TEIV and TEAIV models with a constant death rate over all *E, A* and *I* stages, when cell death was included in the model. Cytotoxic T lymphocytes eliminate virus-infected cells through apoptotic pathways through recognizing certain receptors expressed on the surface of infected cells [99]. Thus, cell death may be better represented as a function of the cell ramping-up budding rate. That is, ramping-up budding state, *i*, should have a *d*_*i*_ death rate that changes with the budding rate (see supplemental material for the full generalized model). This modelling idea was first advanced by Nelson *et al*. [69] for HIV to describe how the cell virus production rate can be coupled with cell age, whereby increased budding leads to more viral surface peptide and therefore to a higher probability of CTL recognition of an infected cell, and a higher rate of infected cell removal. For SARS-CoV-2, CTL-mediated immunity is known to be important [100]. Therefore, in future work we are interested in advancing model fits that more accurately functionally represent infected cell death to further understand the adaptive immune response against SARS-CoV-2; the general TEAIV model with CTL-based cell death ramping is provided in Eq. (S1) with corresponding ℛ_0_ (Eq. (S5)).

Different viral families, e.g., Retroviridae (HIV), Orthomyxoviridae (Influenza), and Coronaviridae (SARS-CoV-2), can exhibit strikingly different infection timescales when TEIV-based models are fit to viral load or cytokine data [101, 102]. These differences can stem not only from broad evolutionary divergence but also from intra-family variation, where distinct viral species, or specific strains, exhibit distinct replication kinetics and immune evasion strategies from other viral strains within the same family. For instance, single mutations affecting entry efficiency, replication rate, or immune evasion can substantially shift viral production and clearance, leading to altered viral fitness and model structure [23]. Therefore, a flexible modelling framework that accommodates variations in eclipse duration and viral budding dynamics is essential. Our current study is the first step in our aim to systematically explore a broad space of mechanistic model formulations, employing consistent fitting approaches, to identify the model(s) that best captures the biological realities of a given pathogen. This would enable inference not only of optimal parameter values, but also of the most plausible mechanistic architecture governing within-host viral dynamics for a specific viral strain.

## 5 Acknowledgements

This work was supported by NSERC, CIHR, AI4PH, and the York University Research Chair program.

## S1 Appendix

### S1.1 Reproduction number derivation for general TEAIV model

Here we derive the reproduction number, ℛ _0_, for the most general TEAIV model with cell death rates, *d*_*i*_, dependent on infectious cell ramping stage. We then simplify to consider ℛ _0_ for the case of no infectious cell death and constant cell death, *d*_*i*_ = *D*, as used for data fitting in the current work.

The general TEAIV model can be formulated as follows:

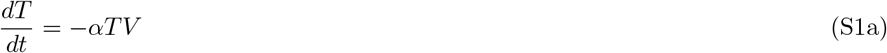

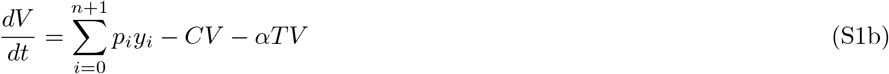

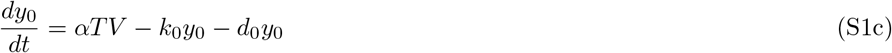

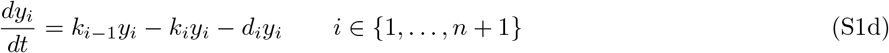

We have, for the main model, *y* = (*E, A*_1_, …, *A*_*n*_, *I*), *p*_0_ = 0, *p*_*n*+1_ = *B*, and *p*_*i*_ *≤ p*_*i*+1_ for 0 *≤ i ≤ n*. For ease of notation and generality, we have added progression out of the final compartment, at a rate *k*_*n*+1_. To keep notation consistent with the main model, *n* counts the number of intermediate stages rather than the number of infected or infectious stages.

Any point in the state space with *V* = 0 and *y*_*i*_ = 0, *i* ∈ {0, …, *n* + 1} is an equilibrium, which we refer to as an infection-free equilibrium. Note that this set of equilibrium solutions is a line in the state space: the *T* -axis. Every point on this line is an equilibrium. Following the trend in epidemic modelling, by stability of infection-free solutions we mean stability to perturbations off this line.

The Jacobian Matrix about an infection-free equilibrium with *T* = *T*_0_ can be partitioned as

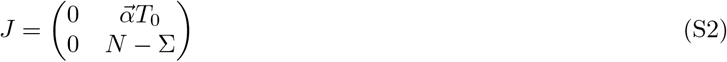

Where 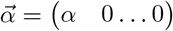

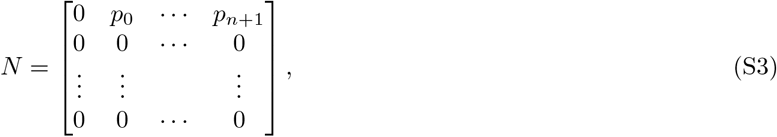

and Σ is a triangular matrix with nonzero entries on the main diagonal and subdiagonal and zero entries everywhere else.

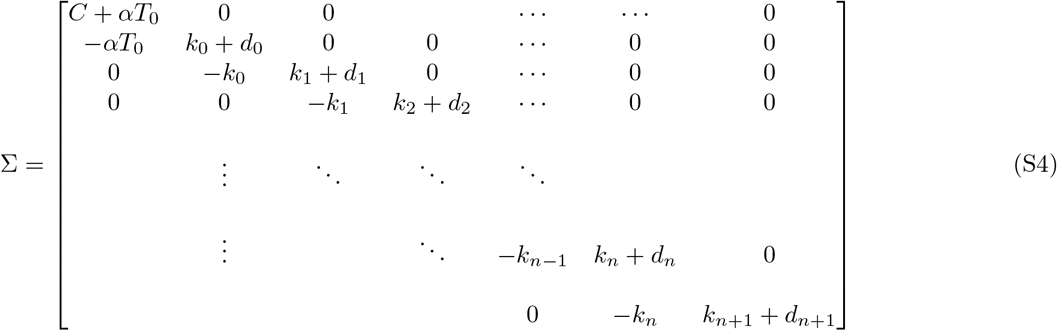

The basic reproduction number can be defined as the spectral radius of the next generation matrix, *N*Σ^*−*1^, and is a threshold for the stability of each equilibrium[103]. Note that the zero eigenvalue of *J* is associated with the line of equilibrium solutions. Perturbations that involve only *T*_0_ are neutrally stable.

Note that *N* is a rank one matrix. This implies the basic reproduction number is

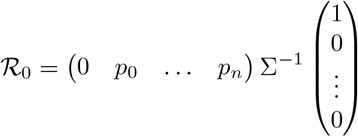

Inverting Σ, we find the following entries in the first column:

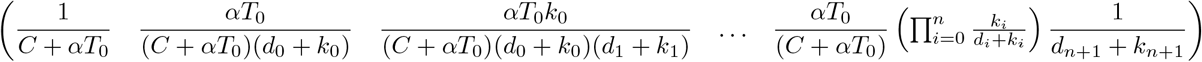

Thus

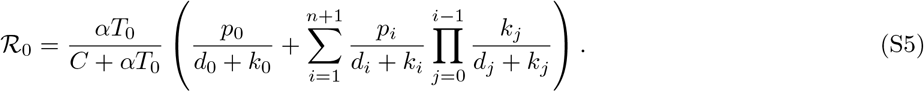

The ratio 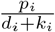 is the expected amount of virus produced by cell in the *i*^th^ stage of infection, and the product inside the summation is the fraction of cells that survive from initial infection to the *i*^th^ stage of infection. Summing these terms over all infected compartments, *i* ∈ {0, …, *n*}, gives the burst size, or the expected number of virions produced over an infected cell’s life. The ratio 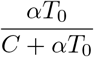 is the fraction of virions that enter and infect susceptible cells. Note that we have used *α* as both the rate cells are infected, and the rate of virus removal during the infection process. More accurately, we could use two paramters here to reflect the possibility of viruses that are bound in this process but do not lead to infection, and to account for the different units of measure between free virus and cells.

An interesting special case is when *p*_0_ = 0, *d*_*i*_ = *D*, and *k*_*i*_ = (*n* + 1)*k, i* ∈ {0, …, *n*}, *k*_*n*+1_ = 0 so that the first stage is a true eclipse stage and the rates in the intermediate stages are identical.

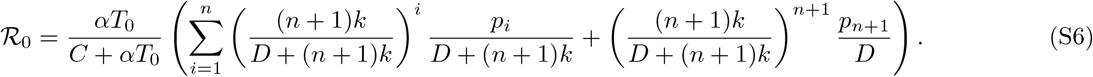

For the special case of arithmetic ramp-up considered in the main manuscript, we have 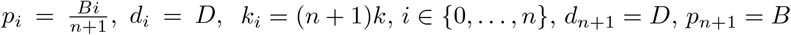, and *k*_*n*+1_ = 0.

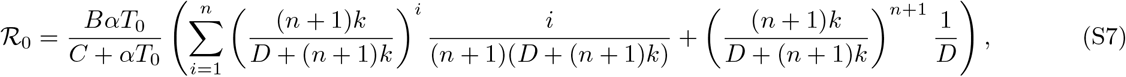

and for the case of no cell death during the budding and eclipse stages we have *d*_*i*_ = 0, *i* ∈ {0, …, *n*} and *d*_*n*+1_ = *D* leading to

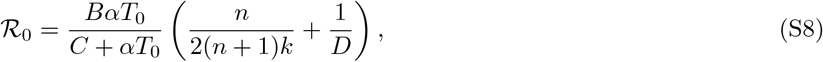

### S1.2 Ramping Functions

Throughout the manuscript, we have taken *n* ramp-up stages, *A*_*i*_, each producing virus at a budding rate *p*_*i*_ with 0 ≤ *p*_*i*_ ≤ *p*_*n*+1_ = *B* so that *B* is the budding rate of a fully infected cell, *I*. We analyzed budding following an arithmetic ramp-up and geometric ramp-up using (3). However, we briefly comment on additional ramp-up considerations that may be useful.

We first note that the geometric ramp-up as written, 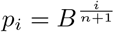 is but one specific choice for a geometric shape parameter. More generally, we require that *p*_*i*_ has units of [*B*] and that *p*_*n*+1_ = *B*. Thus, we could have a more general geometric ramp-up

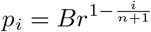

for general shape parameter 0 < *r* < 1 which satisfies the same required properties. We note that the specific case used in (3) is the choice *r* = *B*^*−*1^.

A common saturating process in chemical or biological systems is the use of a Hill function [104, 105] which in the context of budding ramp-up here could take the form

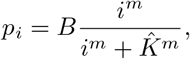

where *m* is the Hill parameter and 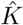 is the half-saturation constant. A limitation of this model in discrete ramp-up is that it does not satisfy the requirement that *p*_*n*+1_ = *B* but rather lim_*i→∞*_ *p*_*i*_ = *B*. Thus, one advantage of the Hill function is in a large *n* limit of virus budding. To take this limit we define

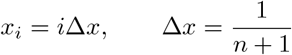

and rewrite the Hill function as

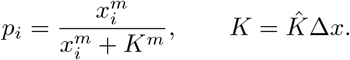

We similarly define a budding-stage density *A*(*x*) and connect it to the compartment *A* _*i*_ via,

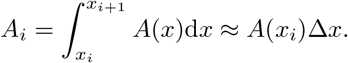

From here we can write the large *n* limit of the sum as

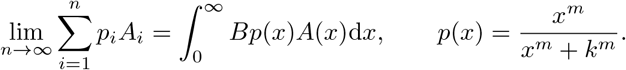

The large *n* continuum limit of (2f) and (2g) then becomes

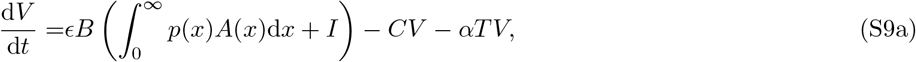

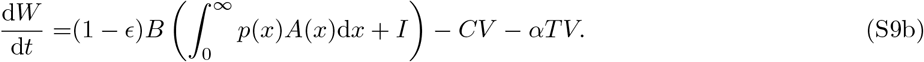

The discrete model has *n* compartments *A*_*i*_ that satisfy

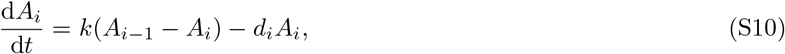

where following Table 1, *k* = *k*_0_(*n*+1) (we take *k*_0_ = 5 throughout the manuscript). Recognizing that *n*+1 = (Δ*x*)^*−*1^ and taking the limit as *n → ∞* allows us to write (S10) as

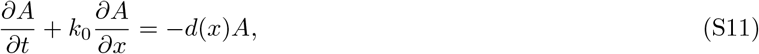

transforming the *n* compartmental ordinary differential equations into a single partial differential equation with a drift term.

### S1.3 Burst size for arithmetic and geometric ramping

**Figure S1:**
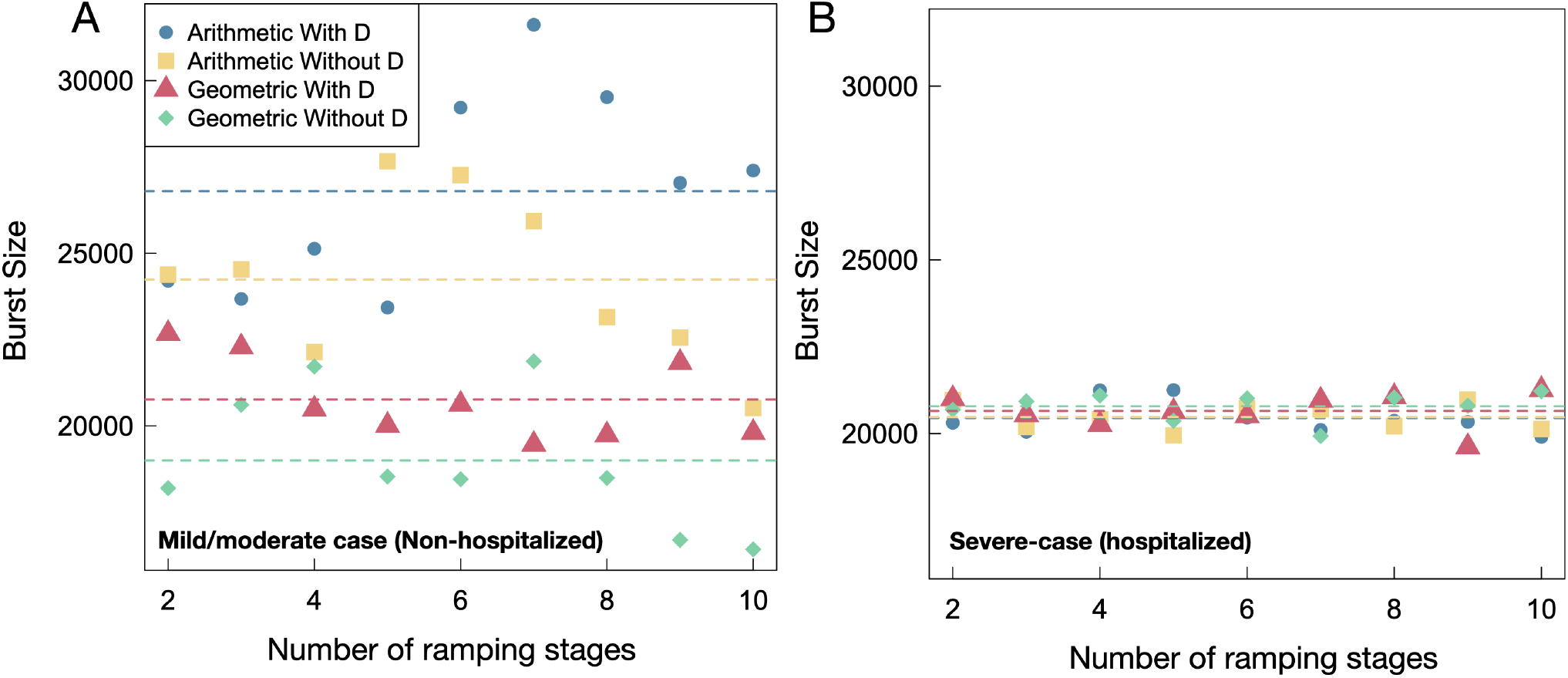
Burst size numerical estimates from Eq. 4 for non-hospitalized (A) and hospitalized (B) model fits. Since burst size is the factor of ℛ_0_ dependent on the ramp up function (Eq. S5), it follows the same trends as *R*_0_ (Fig 4).

### Parameters estimations

**Table S1:**
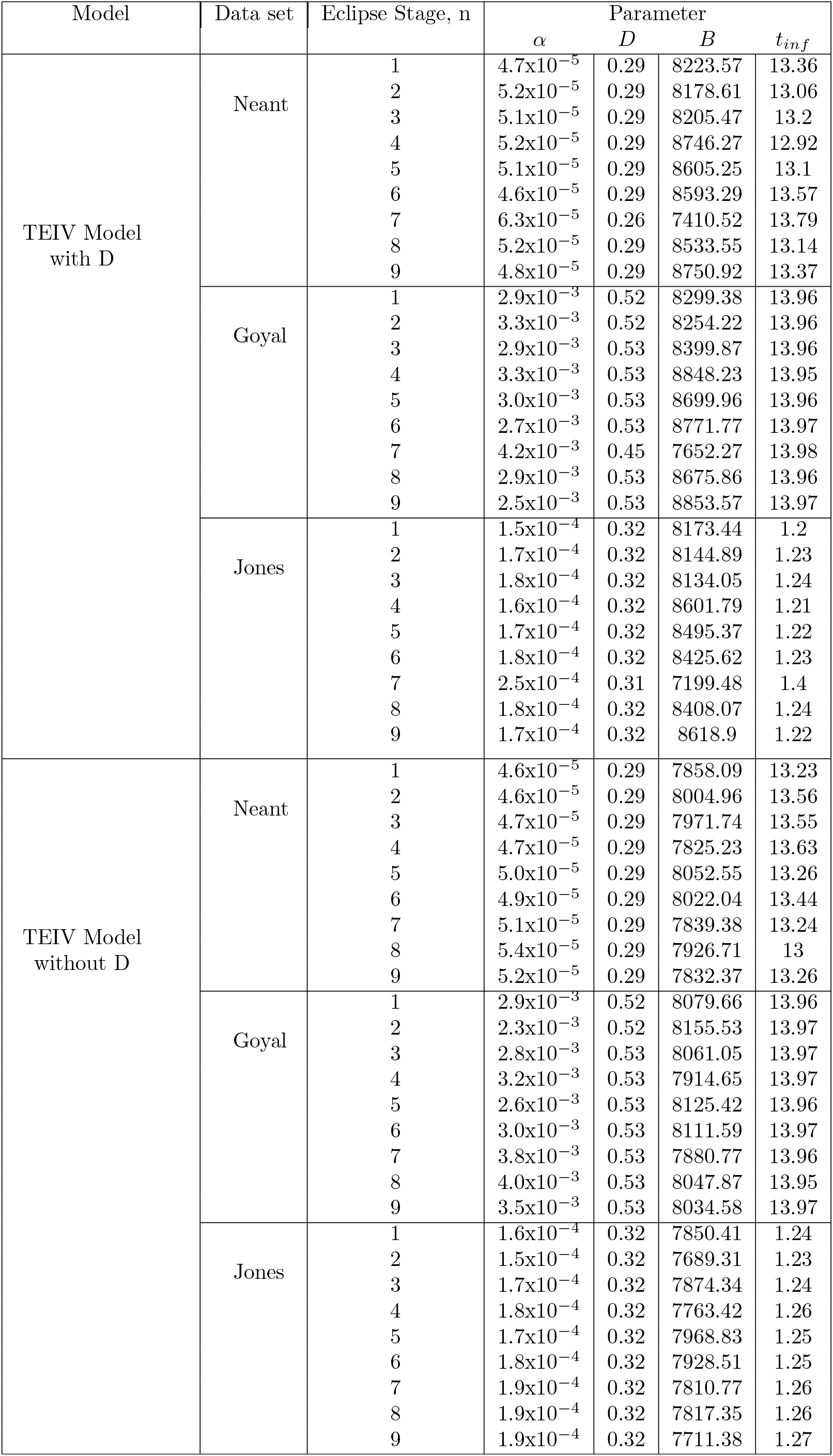
Fitting parameters for all data sets for all investigated *E*_*i*_ Stages for the TEIV model (Eq. 2)

**Table S2:**
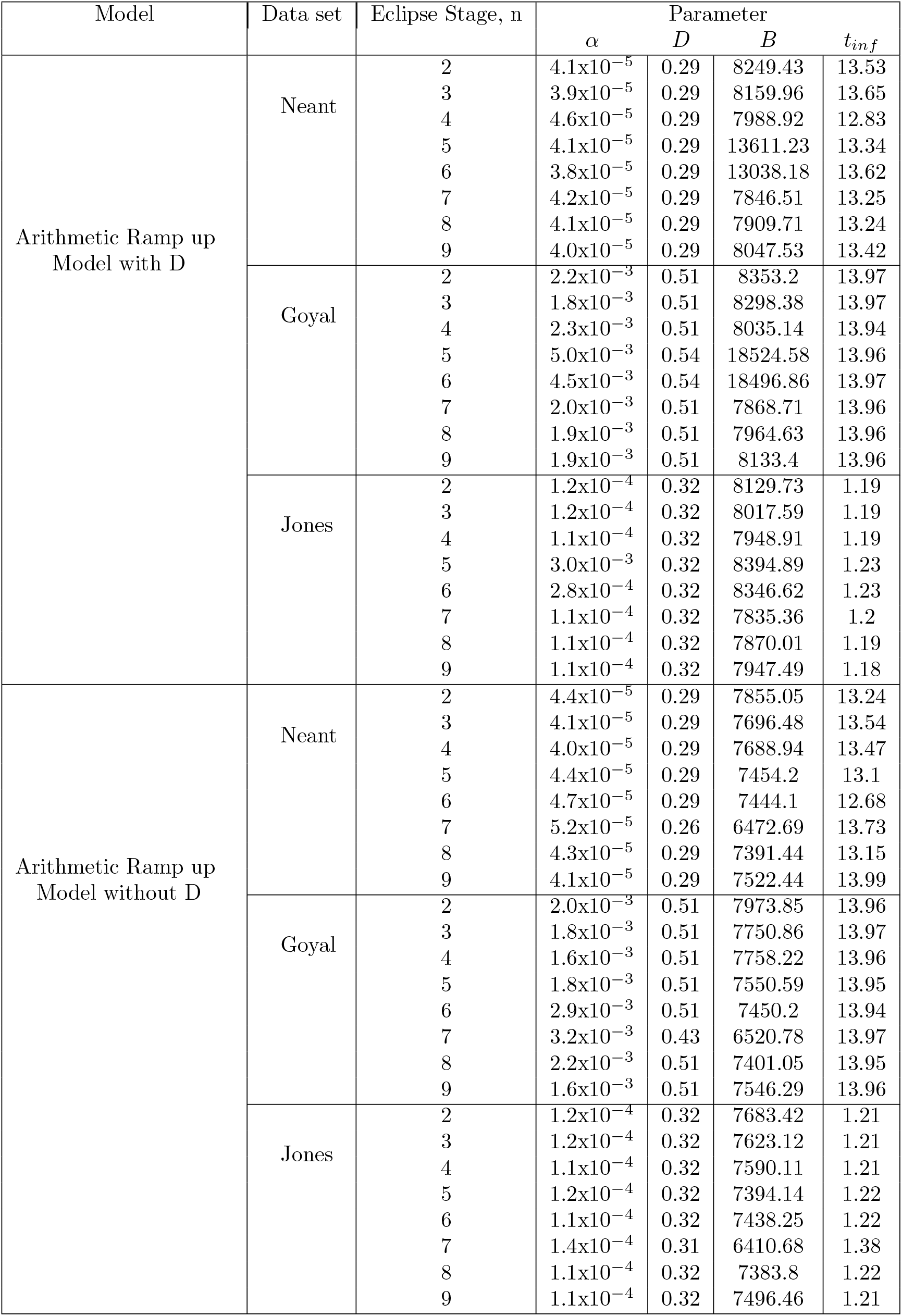
Fitting parameters for all data sets for all investigated *E* + *A*_*i*_ Stages for the TEAIV arithmetic ramp-up model (Eq. 2)

**Table S3:** Fitting parameters for all data sets for all investigated *E* + *A*_*i*_ Stages for the TEAIV geometric ramp-up model (Eq. 2)

| Model | Data set | Eclipse Stage, n | Parameter |  |  |  |
| --- | --- | --- | --- | --- | --- | --- |
| | | | $\alpha$ | $D$ | $B$ | $t_{inf}$ |
| Geometric Ramp up<br>Model with D | Neant | 2 | $4.8 \times 10^{-5}$ | 0.29 | 8435.87 | 13.35 |
| | | 3 | $4.8 \times 10^{-5}$ | 0.29 | 8454.41 | 13.34 |
| | | 4 | $5.1 \times 10^{-5}$ | 0.29 | 8553.7 | 13.04 |
| | | 5 | $4.8 \times 10^{-5}$ | 0.29 | 8336.52 | 13.41 |
| | | 6 | $4.4 \times 10^{-5}$ | 0.29 | 8655.54 | 13.62 |
| | | 7 | $4.8 \times 10^{-5}$ | 0.29 | 8474.69 | 13.32 |
| | | 8 | $4.8 \times 10^{-5}$ | 0.29 | 8453.07 | 13.26 |
| | | 9 | $5.2 \times 10^{-5}$ | 0.29 | 8260.97 | 13.06 |
| | Goyal | 2 | $3.0 \times 10^{-3}$ | 0.52 | 8598.06 | 13.96 |
| | | 3 | $3.0 \times 10^{-3}$ | 0.52 | 8716.32 | 13.97 |
| | | 4 | $2.8 \times 10^{-3}$ | 0.53 | 8600.68 | 13.96 |
| | | 5 | $3.1 \times 10^{-3}$ | 0.53 | 8555.63 | 13.97 |
| | | 6 | $2.7 \times 10^{-3}$ | 0.53 | 8885.8 | 13.97 |
| | | 7 | $2.4 \times 10^{-3}$ | 0.53 | 8606.99 | 13.96 |
| | | 8 | $2.9 \times 10^{-3}$ | 0.53 | 8529.77 | 13.96 |
| | | 9 | $3.2 \times 10^{-3}$ | 0.53 | 8432.74 | 13.96 |
| | Jones | 2 | $1.6 \times 10^{-4}$ | 0.32 | 8295.26 | 1.21 |
| | | 3 | $1.7 \times 10^{-4}$ | 0.32 | 8264.45 | 1.23 |
| | | 4 | $1.6 \times 10^{-4}$ | 0.32 | 8496.18 | 1.21 |
| | | 5 | $1.7 \times 10^{-4}$ | 0.32 | 8204.28 | 1.24 |
| | | 6 | $1.6 \times 10^{-4}$ | 0.32 | 8393.52 | 1.22 |
| | | 7 | $1.7 \times 10^{-4}$ | 0.32 | 8356.8 | 1.23 |
| | | 8 | $1.7 \times 10^{-4}$ | 0.32 | 8378.08 | 1.23 |
| | | 9 | $1.7 \times 10^{-4}$ | 0.32 | 8176.23 | 1.23 |
| Geometric Ramp up<br>Model without D | Neant | 2 | $4.5 \times 10^{-5}$ | 0.29 | 7960.69 | 13.73 |
| | | 3 | $5.0 \times 10^{-5}$ | 0.29 | 7856.07 | 13.26 |
| | | 4 | $4.8 \times 10^{-5}$ | 0.29 | 8003.03 | 13.36 |
| | | 5 | $4.7 \times 10^{-5}$ | 0.29 | 8002.19 | 13.47 |
| | | 6 | $5.0 \times 10^{-5}$ | 0.29 | 7902.22 | 13.26 |
| | | 7 | $4.5 \times 10^{-5}$ | 0.29 | 7906.51 | 13.65 |
| | | 8 | $4.6 \times 10^{-5}$ | 0.29 | 8074.02 | 13.51 |
| | | 9 | $4.6 \times 10^{-5}$ | 0.29 | 7941.21 | 13.58 |
| | Goyal | 2 | $2.7 \times 10^{-3}$ | 0.52 | 8151.05 | 13.98 |
| | | 3 | $3.6 \times 10^{-3}$ | 0.53 | 7881.46 | 13.96 |
| | | 4 | $2.6 \times 10^{-3}$ | 0.53 | 8055.47 | 13.96 |
| | | 5 | $2.8 \times 10^{-3}$ | 0.53 | 8078.21 | 13.97 |
| | | 6 | $3.0 \times 10^{-3}$ | 0.53 | 7974.32 | 13.96 |
| | | 7 | $2.2 \times 10^{-3}$ | 0.53 | 7965.85 | 13.97 |
| | | 8 | $2.8 \times 10^{-3}$ | 0.53 | 8212.5 | 13.97 |
| | | 9 | $2.3 \times 10^{-3}$ | 0.53 | 8119.62 | 13.97 |
| | Jones | 2 | $1.7 \times 10^{-4}$ | 0.32 | 7791.04 | 1.24 |
| | | 3 | $1.7 \times 10^{-4}$ | 0.32 | 7835.65 | 1.24 |
| | | 4 | $1.6 \times 10^{-4}$ | 0.32 | 7938.74 | 1.24 |
| | | 5 | $1.7 \times 10^{-4}$ | 0.32 | 7914.96 | 1.24 |
| | | 6 | $1.7 \times 10^{-4}$ | 0.32 | 7827.08 | 1.25 |
| | | 7 | $1.7 \times 10^{-4}$ | 0.32 | 7849.93 | 1.25 |
| | | 8 | $1.7 \times 10^{-4}$ | 0.32 | 7894.24 | 1.25 |
| | | 9 | $1.7 \times 10^{-4}$ | 0.32 | 7792.68 | 1.26 |

**Table S4:** Estimated Population Parameters from fitting to hospitalized (top - blue header) and non-hospitalized (bottom - orange header) data in Monolix. *B, D*, and *t*_*inf*_ are population estimations of virion budding rate from infected cells, death rate, and time of infection, respectively.

| $E + A_i$ Stages | TEAIV Model | | | | TEIV Model | |
| --- | --- | --- | --- | --- | --- | --- |
| | Arithmetic Ramp-Up | | Geometric Ramp-Up | | w/out $D$<br>$B/D/t_{inf}$ | w/ $D$<br>$B/D/t_{inf}$ |
| | w/out $D$<br>$B/D/t_{inf}$ | w/ $D$<br>$B/D/t_{inf}$ | w/out $D$<br>$B/D/t_{inf}$ | w/ $D$<br>$B/D/t_{inf}$ | | |
| Hospitalized cohort fit parameters |  |  |  |  |  |  |
| 1 | NA | NA | NA | NA | 7855.88/0.36/13.18 | 8223.4/0.36/13.35 |
| 2 | 6569.66/0.32/13.5 | 6778.03/0.32/13.56 | 6817.83/0.33/13.55 | 7389.2/0.33/13.69 | 8005.03/0.36/13.56 | 8178.53/0.37/13.05 |
| 3 | 6504.36/0.33/13.52 | 6675.38/0.32/13.38 | 6892.03/0.33/13.54 | 7223.52/0.33/13.58 | 6924.06/0.33/13.94 | 7264.76/0.33/13.6 |
| 4 | 6369.72/0.32/13.53 | 7062.16/0.32/13.58 | 6943.27/0.33/13.52 | 7119.71/0.33/13.76 | 6729.25/0.33/13.46 | 7439.97/0.33/13.59 |
| 5 | 6219.43/0.32/12.84 | 7057.1/0.32/13.59 | 6699.4/0.33/13.49 | 7245.35/0.33/13.61 | 6614.73/0.33/13.63 | 7234.98/0.33/13.37 |
| 6 | 6654.49/0.33/13.67 | 6787.13/0.32/13.63 | 6908.5/0.33/13.7 | 7201.72/0.33/13.29 | 7002.97/0.33/13.58 | 7312.34/0.33/13.31 |
| 7 | 6447.24/0.32/13.25 | 6666.54/0.32/13.36 | 6549.82/0.33/13.57 | 7355.43/0.33/13.72 | 6636.91/0.33/13.8 | 7471.27/0.33/13.63 |
| 8 | 6290.75/0.32/13.71 | 6751.03/0.32/13.2 | 6702.56/0.32/13.56 | 7389.2/0.33/13.69 | 6763.31/0.33/13.8 | 7162.81/0.33/13.2 |
| 9 | 6526.36/0.32/13.27 | 6738.36/0.32/13.63 | 6832.92/0.33/13.77 | 6881.61/0.33/13.72 | 6829.15/0.32/13.89 | 7528.92/0.33/13.81 |
| 10 | 6261.33/0.32/13.16 | 6593.39/0.32/13.53 | 6968.95/0.33/13.78 | 7457.12/0.33/13.63 | 6898.87/0.33/13.58 | 7260.1/0.33/13.46 |
| Non-hospitalized cohort fit parameters |  |  |  |  |  |  |
| 1 | NA | NA | NA | NA | 16549.68/0.75/10.96 | 22657.53/0.76/10.16 |
| 2 | 17453.76/0.75/10.31 | 20286.99/0.76/10.14 | 13076.47/0.72/12.2 | 19348.98/0.74/10.99 | 14269.28/0.76/11.52 | 19346.28/0.75/10.92 |
| 3 | 17674.71/0.76/10.07 | 19735.59/0.76/10.07 | 15197.9/0.74/11.53 | 19297.3/0.75/11.01 | 13039.85/0.74/12.14 | 14895.4/0.74/12.02 |
| 4 | 15887.85/0.76/10.3 | 20876.45/0.76/9.89 | 15994.3/0.74/11.38 | 17719.12/0.75/11.3 | 14937.24/0.75/11.53 | 16130.21/0.74/11.85 |
| 5 | 20046.43/0.77/9.63 | 19416.87/0.76/10.02 | 13630.62/0.74/11.94 | 17037.6/0.74/11.53 | 14116.35/0.74/11.85 | 17337.01/0.74/11.57 |
| 6 | 19718.79/0.77/9.7 | 24521.28/0.77/9.49 | 13378.31/0.73/12.01 | 17543.4/0.74/11.44 | 13132.23/0.73/12.27 | 16226.73/0.73/11.95 |
| 7 | 18727.96/0.77/9.73 | 26500.35/0.77/9.27 | 16061.26/0.74/11.31 | 16804.6/0.75/11.5 | 12853.66/0.74/12.23 | 16031.5/0.74/11.89 |
| 8 | 16500.72/0.76/10.03 | 24719.57/0.77/9.4 | 13385.6/0.73/12.06 | 16515.6/0.73/11.7 | 13957.79/0.74/11.96 | 15767.34/0.74/12.01 |
| 9 | 16060.36/0.76/10.07 | 22306.1/0.76/9.58 | 12242.8/0.74/12.31 | 18547.19/0.74/11.24 | 12504.65/0.74/12.4 | 16970.03/0.73/11.8 |
| 10 | 14417.45/0.75/10.45 | 23228.95/0.78/9.37 | 12197.9/0.75/12.15 | 16817.69/0.74/11.61 | 12454.21/0.73/12.54 | 16096.3/0.74/11.96 |

## Footnotes

1 There is a general class of functions that could be used for geometric ramp-up as discussed in Section S1.2 in the Supplementary Material.

